# Are place cells just memory cells? Memory compression leads to spatial tuning and history dependence

**DOI:** 10.1101/624239

**Authors:** Marcus K. Benna, Stefano Fusi

**Author notes:** M.K.B. and S.F contributed equally to this work.

## Abstract

The observation of place cells has suggested that the hippocampus plays a special role in encoding spatial information. However, place cell responses are modulated by several non-spatial variables, and reported to be rather unstable. Here we propose a memory model of the hippocampus that provides a novel interpretation of place cells consistent with these observations. We hypothesize that the hippocampus is a memory device that takes advantage of the correlations between sensory experiences to generate compressed representations of the episodes that are stored in memory. A simple neural network model that can efficiently compress information naturally produces place cells that are similar to those observed in experiments. It predicts that the activity of these cells is variable and that the fluctuations of the place fields encode information about the recent history of sensory experiences. Place cells may simply be a consequence of a memory compression process implemented in the hippocampus.

**Significance Statement:** Numerous studies on humans revealed the importance of the hippocampus in memory formation. The rodent literature instead focused on the spatial representations that are observed in navigation experiments. Here we propose a simple model of the hippocampus that reconciles the main findings of the human and rodent studies. The model assumes that the hippocampus is a memory system that generates compressed representations of sensory experiences using previously acquired knowledge about the statistics of the world. These experiences can then be memorized more efficiently. The sensory experiences during the exploration of an environment, when compressed by the hippocampus, lead naturally to spatial representations similar to those observed in rodent studies and to the emergence of place cells.

## Introduction

Several studies show that neurons in the hippocampus encode the position of the animal in its environment and as a consequence they have been named “place cells” (see e.g. (1)). Here we propose a novel interpretation of the observation of place cells by suggesting that their response properties actually reflect a process of memory compression in which the hippocampus plays a fundamental role. Our interpretation contributes to the reconciliation between two dominant, but apparently different points of view: one involving the hippocampus in spatial cognitive maps and navigation, versus another one that considers the hippocampus playing a broad role in episodic and declarative memory (see e.g. (2, 3)).

The first view is supported by the observation of place cells, some of which exhibit responses that are easily interpretable as the cells tend to fire only when the animal is in one particular location (single field place cells). However, it is becoming clear that in many brain areas, including the hippocampus and entorhinal cortex, neural responses are very diverse (4–8), highly variable in time (9, 10) and modulated by multiple variables (8, 11–13). Place cells might respond at single or multiple locations, and multiple visitations of the same location typically elicit different responses. Part of this diversity can be explained by assuming that each neuron responds non-linearly to a different combination of multiple external or internal variables (mixed selectivity; see e.g. (5, 6)). The variability might be due to the fact that some of these variables are not being monitored in the experiment, and hence contribute to what appears to be noise. Some of the components of the variability probably depend on the variables that are represented at the current time, but some others might depend also on the recent history, or in other words, they might be affected by the storage of recent memories.

The model of the hippocampus we propose not only predicts that place cells should exhibit history effects, but also that their spatial tuning properties simply reflect an efficient strategy for storing correlated patterns. Much of the theoretical work on the memory capacity of neural networks is based on the assumption that the patterns representing memories are random and uncorrelated (see e.g. (14–18)). This assumption is not unreasonable for long-term storage, despite the fact that most of our sensory experiences are highly correlated. Indeed, to efficiently store correlated episodes, it is desirable to preprocess the new memories and compress them before they are placed in long-term memory. Ideally, one would want to extract the uncorrelated (and hence incompressible) components of the new input patterns and store only those. However, this form of preprocessing and compression, which would lead to uncorrelated patterns of synaptic modifications, has not been explicitly modeled in previous theoretical studies of memory capacity.

We hypothesize that this preprocessing of the incoming stream of potential episodic memories is to some extent carried out in the hippocampus, which in itself has long been considered as a memory storage system, albeit with a longest time constant that is shorter than that of cortex (19, 20). New sensory episodes are often similar to old ones and hence their neural representations are likely to be correlated to those of sensory experiences that are already in memory. Storing directly the neural representations of these sensory episodes would be inefficient because they are redundant. However, since the hippocampus is highly structured and contains distinct regions that might implement different stages of processing, it can transform and compress incoming sensory information with respect to other memories already stored there. An important role of the hippocampus in compressing memories has already been suggested in (21) (see also the Discussion).

We have built a concrete neural network model that implements this compression process in a manner broadly consistent with the known neuroanatomy. The model consists of a three-layer network in which the first layer represents the sensory inputs (which would already have been processed by the sensory areas corresponding to different modalities). The feed-forward synapses that connect it to the second layer are continuously modified to create a compressed, sparse representation of the inputs. This layer implements a form of statistical learning, as the compressed representations are based on the statistics of recent sensory experiences (and possibly input from cortical long-term memory in the case of familiar environments, as discussed below). A third layer is used to store the memories, i.e., specific episodes. This architecture and the computational principles are similar to those proposed in (22) (see also the Discussion). We show that the plasticity of the feed-forward synapses robustly improves the memory capacity compared to a network of the same architecture, but with fixed feed-forward weights (which would implement a random projection of the correlated patterns).

Furthermore, the model explains quite naturally the emergence of place cells in the second and the third layer, without the need to assume any special role of physical space compared to other external variables. Compressing sensory inputs of an animal in a given environment automatically leads to the emergence of cells whose activity is strongly modulated by its position in the environment, because many experiences of the animal depend on its position, and the latter is thus highly informative about the former. By the same token, processing sensory inputs correlated with other external variables would encourage cells to develop receptive fields in the space of those variables (23), since the computational principle of compressing inputs for efficient storage is agnostic to the nature of the variables that induce the correlations.

Moreover, such models with ongoing plasticity predict that the neural representation encoding a sensory episode will differ depending on the previous experiences of the animal, that is, it will be history-dependent. In particular, synaptic weights are constantly modified (and correspondingly the neural tuning properties change) in order to update the statistical model of the environment and to store new sensory experiences (episodes) in memory. The resulting place cell responses can be modulated by any variable that describes relevant aspects of the experienced sensory inputs. We will show in simulations that even in the absence of novel salient events such as the delivery of a reward, the place fields constantly fluctuate to reflect the most recent changes in the input statistics, though they remain sufficiently stable to decode position.

## Results

### Storing correlated patterns efficiently

Most of the patterns we would like to store in memory are likely to be highly correlated, as our experiences are often similar to each other. Storing correlated patterns in artificial neural networks typically requires a synaptic learning rule that is more complicated than simple Hebbian plasticity to avoid a bias towards the memory components shared my multiple memories. Simple extensions such as the perceptron learning rule can already deal with many forms of correlations. However, storing correlated patterns in their original format is rarely the optimal strategy and often it is possible to greatly increase the memory capacity by constructing compressed neural representations that explicitly take into account the correlations. This form of preprocessing can be illustrated with a simple example in which the patterns to be memorized are organized as in an ultrametric tree (see e.g. (24) and Figure 1a). To generate these correlated patterns one starts from *p* uncorrelated random patterns (the ancestors at the top of the tree). In these patterns each neuron is either active or inactive with equal probability, as in the case of the Hopfield model (14). One can then generate *k* descendants for each ancestor by resampling randomly with a certain probability 1 – *γ* the activation state of each of the neurons in the patterns representing the ancestors. The parameter *k* is called the branching ratio of the tree. The total number of descendants is *p k*, and these are the patterns that we intend to store in memory. The descendants that come from the same ancestor will be correlated, since they will all be similar to their ancestor. This is a very basic scheme for generating correlations between patterns, which has been studied to extend the Hopfield model to the case of correlated attractors (25–27).

**Fig. 1.**
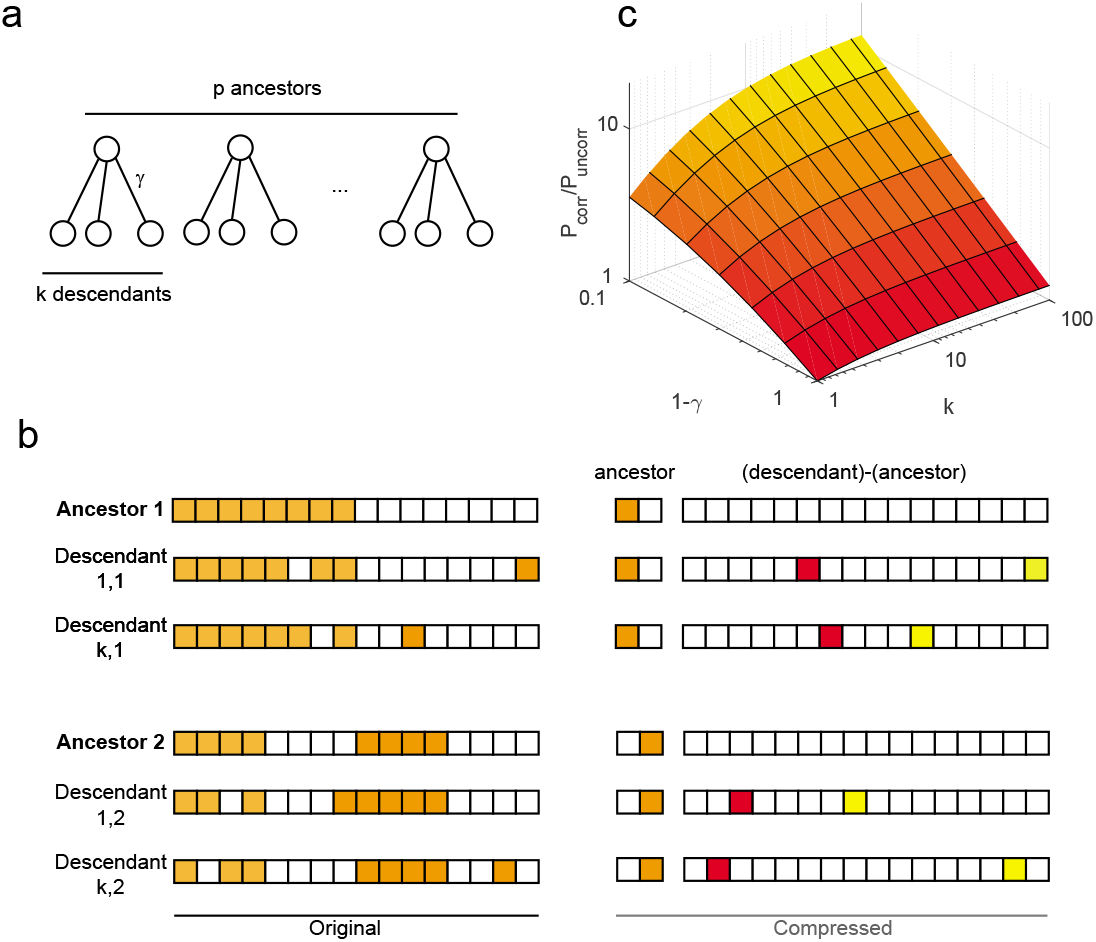
Storing efficiently correlated patterns in memory. (a): Schematic of an ultra-metric tree with *p* ancestors and *k* descendants per ancestor used to generate correlated patterns. (b): A possible scheme to take advantage of the correlations and generate compressed representations that are sparse and hence more efficently storable. (c): Total number *P_corr_* of correlated patterns generated from a tree model with parameters *p, k* and *γ* that can be stored using a simple compression strategy, divided by the number of patterns *P_uncorr_* that could be stored (using approximately the same number of neurons and synapses) if the patterns were uncorrelated. Therefore, the plot shows the relative advantage of using a compression strategy compared to storing incompressible patterns as a function of *k* and *γ*.

One simple strategy to efficiently store these patterns in memory is to store the ancestors in one network, and the differences between the ancestors and their descendants in another network (see Figure 1b). These differences are approximately uncorrelated, and they can be significantly sparser than the original patterns of activity (for *γ* close to 1; see also (28)). Indeed, most of the neurons have the same level of activation in the ancestors and in its descendants, and hence the difference is zero. A sparse representation is one in which only a relatively small fraction *f* of the neurons is active. Sparse random patterns contain less information than dense patterns (the information per neuron scales approximately as *f*, which is called the coding level, see e.g. (29)). However, the number of random sparse patterns that can be stored in memory can be much larger than the number of storable dense patterns (i.e., patterns with *f* = 1/2) (16–18, 29–32), mostly because the interference between memories is strongly reduced. In this simple scheme, which first extracts and stores the ancestors, it is possible to compress the information about the descendants simply by constructing sparse representations that take into account the already acquired information about the ancestors. Even though the amount of information per pattern is smaller for sparser representations, there is no loss of information, because storing differences between ancestors and descendants actually requires fewer bits than storing the full patterns representing the descendants.

The relative advantage of this scheme as measured by the improvement factor of the total number of retrievable descendant patterns compared to storing uncorrelated patterns is estimated in the Methods and summarized in Figure 1c. As *k* increases, more of the patterns to be stored become correlated and their overall compressibility increases. The improvement factor also increases, because the scheme that we discussed can take advantage of the increased compressibility. From the formula in the Methods, it is clear that the improvement increases approximately linearly with *k*, when *γ* is fixed close to 1. The improvement also increases as *γ* tends to 1, i.e., when the descendants become more similar to their common ancestor and hence the patterns are more compressible.

### Generating compressed representations using sparse auto-encoders

The scheme we just discussed illustrates a simple strategy for taking advantage of the correlations between the patterns to be memorized. Essentially, eliminating correlations allows us to increase the level of sparseness, which in turn leads to a larger memory capacity. However, it is unclear whether this strategy can actually be implemented in a simple neural network, and more importantly whether it can be extended to real world memories, which are certainly more structured than the ultrametric ones that we considered in the previous section. Here we show that it is possible to construct a simple network that is able to generate compressed representations for arbitrary types of patterns to be memorized. These representations share some of the properties of those discussed in Figure 1 when ultrametric memory patterns are considered.

The network illustrated in Figure 2a comprises two of the three layers of the hippocampal model that we described in the Introduction. The first, input layer, which could be mapped onto entorhinal cortex (EC), encodes sensory experiences and the second layer (possibly the dentate gyrus, DG) their compressed representations. The third layer (CA3) would store specific episodes (i.e., individual input patterns that represent instantaneous sensory experiences), but in this first part of the article we will not simulate it, as we will focus on the geometrical properties of the compressed representations. These representations are not constructed by hand, as in Figure 1, but by using the strategy of sparse auto-encoders: the first two layers are complemented by a reconstruction layer, and the synaptic weights are modified to ensure that the input is faithfully reproduced in the reconstruction layer. We used an algorithm similar to the one introduced by Olshausen and Field to reproduce the neural representations of the visual cortex (33, 34). Recent extensions of this algorithm apply to several important computational problems (35). The main idea is to modify the synaptic weights from the input layer to the second one to build sparse neural representations of the inputs. The weights are chosen to minimize the reconstruction error (of the inputs) when one reads out these second layer representations. The representations obtained using this approach are compressed because of the sparseness that is imposed on them by the algorithm. It is important to stress that the reconstruction layer is used only to determine the weights between the input layer and the layer with the compressed representations. The reconstruction layer presumably does not exist in the brain and indeed other algorithmic approaches do not need it (see e.g. (36–38)). Moreover, in the brain the compressed representations are typically used to perform tasks that involve more than simple memory retrieval or reconstruction. It is only for simplicity and to avoid further assumptions that here we build compressed representations by considering the reconstruction problem (which is analogous to the approach of Olshausen and Field for the visual system; see also the Discussion).

**Fig. 2.**
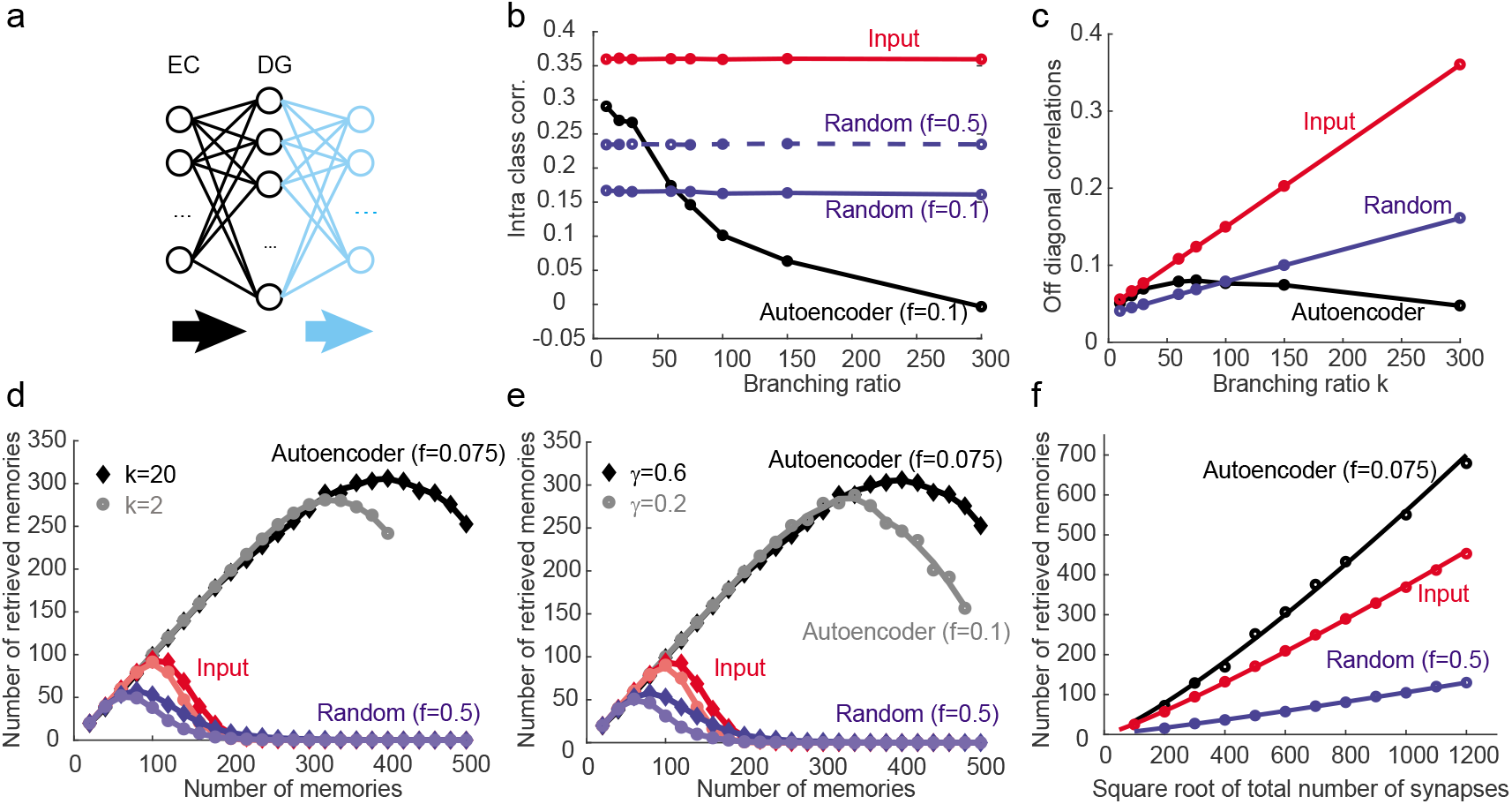
(a): Scheme of the simulated auto-encoder. The input layer (300 neurons, probably mappable to EC), projects to an intermediate layer (DG, 600 neurons). The light blue neurons are introduced to determine the weights from the input to the intermediate layer (they are chosen so that the light blue neurons reproduce the input). (b): Geometry of the compressed neural representations: correlations between the representations of patterns that belong to different descendants of the same ancestor for the inputs (red), the auto-encoder (intermediate layer in panel a; black), and a random encoder (blue) as a function of the branching ratio when the total number of patterns is kept constant (and hence the number of ancestors varies). As *γ* is fixed (*γ* = 0.6), the correlations of the inputs and the random encoder are constant (for the inputs they are *γ*^2^ = 0.36). For the auto-encoder they decrease with the branching ratio *k*. As *k* increases, the environment becomes more compressible and more resources are devoted to the differences between descendants and ancestors, which are approximately uncorrelated. (c): Average of the modulus of the correlations between all patterns as a function of the branching ratio. As the branching ratio increases starting from a small *k*, the representations become more correlated. The representations of the inputs are more correlated than the representations of the auto-encoder and the representations of the random encoder. For high compressibility (large *k*), the auto-encoder has the most decorrelated representations. (d,e): Memory performance of the auto-encoder compared to a random encoder and a readout of the input: the number of retrieved memories is plotted as a function of the total number of memory patterns (changing the number of ancestors). For the auto-encoder we show two curves in (d) that correspond to different branching ratios (*k* = 2, 20) but same *γ* (= 0.6) and two curves in (e) which correspond to different values of *γ* (0.6 and 0.2), the similarity between ancestors and their descendants. As the number of ancestors increases, the quality of retrieval decreases, and the number of correctly retrieved memories reaches a maximum. The auto-encoder outperforms the input and the random encoder. In both (d) and (e) the auto-encoder performs better when the memories are more compressible. (f): memory capacity as a function of the square root of the total number of synapses for auto-encoder, random and input representations. The auto-encoder representations outperform all the others despite the fact that they require four times more synapses when compared to the case in which the inputs are read out directly.

We can now analyze the geometry of these representations and in particular focus on the correlations between the compressed patterns. Indeed, given the simple ultrametric organization of the input patterns, the geometry of the representations is completely characterized by some measure of the similarity between patterns that correspond to descendants of the same ancestor, and between patterns that are descendants of different ancestors. We will compare these correlations to those that characterize the compressed representations obtained from the second layer of the trained auto-encoder. We will consider also the representations obtained in a network in which the neurons in the second layer are randomly connected to the input neurons (random encoder, see e.g. (39–41)). In this network there is no learning; the random weights are chosen when the network is initialized and then they are frozen. The number of neurons of the representations are chosen to be the same as in the auto-encoder to make the comparison of the performance with the auto-encoder as fair as possible.

In Figure 2b we plotted the correlations between the descendants of the same ancestor as a function of the branching ratio when the total number of patterns is kept constant (by varying the number of ancestors). For the inputs and the random encoder they are constant. Note that for the random encoder we show two curves: one for the same coding level *f* = 0.1 as the auto-encoder, and one for the coding level that maximizes the memory capacity of the random encoder (*f* = 0.5, see below). For the auto-encoder the correlations decrease with the branching ratio *k*. This is expected from the abstract scheme that we described above: as *k* increases, the number of ancestors decreases, the number of correlated inputs increases and the full set of patterns to be memorized (the “environment”) becomes more compressible. More memory resources (i.e., plastic synapses that are modified) are devoted to the differences between descendants and ancestors, which are approximately uncorrelated. Indeed, the correlations between the difference patterns are (1 – *γ*)/2, and when the descendants are sufficiently similar to their ancestor (*γ* → 1), these correlations are small.

In Figure 2c we show the average correlations between all patterns as a function of the branching ratio. These are the correlations that would be measured in an experiment when all the recorded patterns of activity are considered. We computed the average of the modulus of the correlations because the average of positive and negative correlations could be close to zero, which might be misinterpreted to mean that the patterns were not correlated. The representations of the inputs are more correlated than the representations of the auto-encoder and those of the random encoder. As the branching ratio increases, the representations become more correlated for the inputs and the random encoder, but this monotonic relationship doesn’t hold for the trained auto-encoder. In fact, for sufficiently large *k* the auto-encoder representations become progressively more decorrelated as the set of memories to be stored becomes more compressible. If the auto-encoder layer is interpreted as the dentate gyrus (DG), this observation would be consistent with the notion that DG is involved in pattern separation (42–45). Interestingly, the model would predict that the representations in DG should become less correlated if the environment is more compressible (see also the Discussion).

We then estimated the memory performance for the autoencoder, for the random encoder, and for reconstruction directly from the input representations, without an intermediate layer. In this case, we kept the branching ratio fixed and we increased the number of ancestors, so that the total number of memories increases. We then estimated the number of memories that were correctly retrieved. A memory was considered correctly retrieved when it was possible to reconstruct the stored memory in the reconstruction layer (i.e., the light blue layer of Figure 2a) with an overlap of at least 0.9 with the original memory. A noisy cue was imposed on the input layer to trigger the memory retrieval process (the noise level of the cue was chosen to be large enough that the average overlap with the original memory was only 0.8). To evaluate the memory performance without an intermediate layer, we connected the input layer directly to the reconstruction layer (see Methods for more details).

As the total number of stored memories increases, the number of retrieved memories also increases until at some capacity level it starts to smoothly decrease. The memory capacity is then defined as the maximum number of memories that can be correctly retrieved. The auto-encoder was trained with sigmoidal activation functions as described above, but we now use a binarized version of the compressed representations to make the comparison of the memory performance with the random encoder and the input as fair as possible. The memory performance is significantly higher when binarization is not imposed.

In Figure 2d we show the number of retrieved memories for the auto-encoder, the input and the random encoder representations. The memory performance is significantly larger for the auto-encoder than for the other two cases, and it is higher for a larger branching ratio (*k* = 20), which corresponds to a more compressible environment. For each curve we used the optimal coding level. The performance of the random encoder is the worst because of the elevated level of noise. It is significantly better, but still much worse than for the auto-encoder, for smaller levels of noise. In Figure 2e we show again the number of retrieved memories in the case in which the branching ratio is always the same, but the similarity *γ* between the descendants and their ancestors varies. The best performance is achieved for the auto-encoder in the case in which the descendants are most similar to their ancestor, which is again the case in which memories are most compressible.

Finally we estimated the memory capacity as a function of the total number of synapses in the system. We showed that the memory capacity is significantly better for auto-encoder representations than for the systems that read out the inputs directly. However, the auto-encoder requires an intermediate layer, and hence a larger number of synapses. In Figure 2e we computed the memory capacity, as in panels d,e, for networks of different sizes. We progressively increased the size of the input (*N* neurons) and we scaled accordingly the number of neurons in the auto-encoder intermediate layer (it was always 2*N*). The total number of synapses was then *N*^2^ in the case in which we directly read out the input, and 4*N*^2^ for the autoencoder. In Figure 2 we show the memory capacity as a function of the square root of the number of synapses for the autoencoder, the random and the input representations. The coding level was chosen to optimize the capacity in the case of *N* = 300. The auto-encoder strategy outperforms all the others and the improvement increases with the size of the network. This analysis shows that the auto-encoder representations can be legitimately called “compressed representations” because they allow for the same performance with a smaller number of synapses.

### Compressing sensory inputs experienced during navigation

We now consider the specific case of navigation. In this case the sensory experiences of an animal that visits the same location multiple times will be different, but still correlated (Figure 3a). Similarly to the case of ultrametric memory patterns described in the previous section, it is possible to take advantage of these correlations to compress the information that is stored in memory. We hypothesize that the hippocampus is involved in this process of compression which leads to sparse, compressed representations of the sensory experiences of the animal during the exploration of an environment. We now present a simple model to illustrate how the compressed representations can be generated and then we show that using these representations leads to a more efficient storage of correlated memories. As in the previous section, for simplicity we construct the compressed representations using a sparse auto-encoder. It is likely that in a more realistic situation in which the animal has to perform a task, the representations are not just shaped by the desire to reconstruct inputs, but also affected by other factors (see the Discussion).

**Fig. 3.**
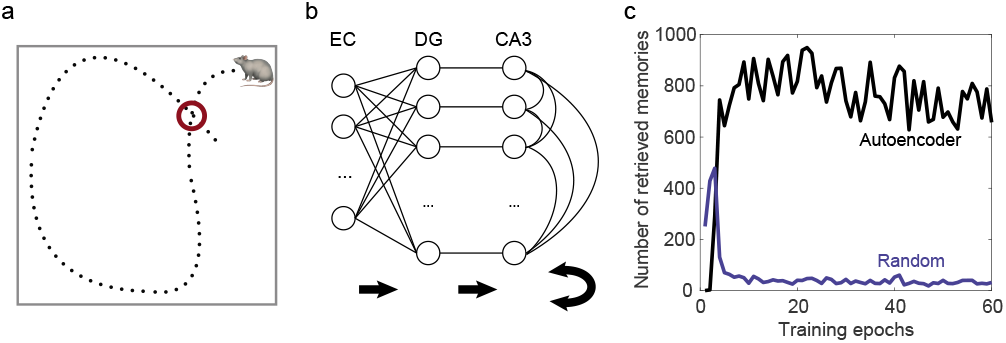
(a): Schematic of a rodent exploring an open field arena. Whenever the animal returns to the same location, its sensory inputs will have some similarity with those experienced during previous visitations of that location. (b): Schematic of the architecture of the network with potential mapping of the layers onto entorhinal cortex and hippocampus. (c): The memory capacity (the number of stored inputs that can be recalled from noisy cues of overlap < 0.7, such that the pattern overlap of the retrieved representation with the original stored pattern is larger than 0.8) as a function of the number of training sessions (exposures to the environment). This illustrates the computational advantage of using even a simple compression algorithm with one layer of learned weights as implemented in our network (black), compared to a network of the same architecture (and coding levels), but with fixed random feed-forward connections (blue).

We assume that the animal is exploring an environment enclosed by four walls which has the shape of a square. The animal can use sensory cues (visual, tactile, olfactory) to determine its position when it is very close to one of the walls and, for simplicity, we assume that it explores the environment by walking along a straight line in a random direction until it reaches another wall. It then repeats this procedure by picking another direction and walking again in a straight line. We also hypothesize that the animal performs a simple form of path integration and hence it knows approximately the distance from the last wall it visited. Finally, we assume that the animal knows the direction of movement by using distal cues. These assumptions would be compatible with the observations that head direction is encoded in entorhinal cortex, one of the major cortical inputs to the hippocampus. Furthermore, we know that the estimate of position decoded from entorhinal cortex (EC) has an accuracy that decreases with the distance from the last visited wall (46), indicating that some form of path integration reset is performed when the animal gets close to a wall.

The model that we simulate is the simple feed-forward network with three layers already described in the Introduction (see also Figure 3b). In the specific case of navigation, the first layer (EC) represents the input, and in our simple example encodes 1) the direction of movement of the animal (47), which could be estimated by looking at distal cues, 2) the distance from the last wall visited (48), and 3) the position along the last wall visited (49) (i.e., the initial position before the animal initiates its excursion to explore the environment). These variables are mixed non-linearly through a random projection to obtain the putative EC representations we use as inputs to our network.

The second layer (DG) contains the compressed representations of the sensory experiences. These representations are learned as described in the previous section by introducing an artificial reconstruction layer (the light blue layer in Figure 2a, not shown in Figure 3b for simplicity) and by imposing that its representations reproduce the inputs.

Finally, the third layer (CA3), is an auto-associative memory system where the memories are stored by modifying the recurrent connections between neurons. The purpose of this layer in our simulations is to obtain a reasonable estimate of the memory capacity beyond simple feed-forward reconstruction (as discussed in the previous section), by incorporating recurrent retrieval dynamics.

The recurrent synaptic weights are modified according to a simple covariance rule, similar to the Hopfield rule (14), to create stable fixed points corresponding to the patterns of activity imposed by the inputs from the second layer. For simplicity we assumed that the number of neurons in the third layer is equal to the number of neurons in the second one and that the third layer basically just copies the compressed sparse representations prepared in the previous layer. This is not entirely unrealistic, since CA3 pyramidal cells do receive only a small number of strong synapses from the dentate gyrus (50). A more realistic third layer with a different number of neurons and with random incoming connections from the second layer would work in a similar fashion. While the second layer neurons are continuous-valued (and typically exhibit a bimodal activity distribution in our simulations), we threshold their activity to obtain binary neural representations suitable for storage in an auto-associative network with binary neurons.

We simulated an animal exploring an unfamiliar environment by discretizing time and computing at every time step the inputs for the current position and direction of motion of the animal by a random projection of the variables described above. These input patterns represent the memories that we intend to store. The patterns are used to determine the compressed representations, which in turn are passed to the third layer, where they are stored in the auto-associative network.

We consider a situation in which the animal explores for several hundred (straight line) trajectories crossing the environment, which is similar to the typical situations studied in experiments on rodent navigation. We compared the performance of the proposed memory system to the performance of an analogous network with the same architecture and coding level in which the input layer is connected to the second layer with fixed random connections. To quantify this memory performance, we count the number of patterns that can be retrieved with sufficiently high fidelity, namely with a pattern overlap of at least 0.8, from noisy cues with an initial overlap less than 0.7 (with the binary pattern that was stored in the auto-associative network). These memory patterns correspond to the sensory inputs experienced by the animal during the preceding exploration of the environment. The same input patterns were used to also train the auto-encoder. The results are reported in Figure 3c, which shows that the performance of the proposed network is substantially better than that of the network with random connections. Note that the coding level in the random network is kept equal to the coding level in the auto-encoder to make the memory performance comparison as fair as possible. The coding level of the auto-encoder changes during the course of learning as the cost function trades off the reconstruction loss against the sparsity penalty. In particular, it initially increases rapidly, which explains the transient increase of the capacity of the random network (which however occurs at a coding level too small to reconstruct the input representations). Once the coding level has approximately stabilized the memory capacity plateaus, but nevertheless the capacity keeps fluctuating between different training epochs, largely due to the changing input statistics for different exposures to the environment (as described below).

### Single neuron properties: the emergence of place cells

Now that we have established that the compressed neural representations allow for a better memory performance, it is interesting to inspect the neural representations obtained in the second and third layers of the network. These are the representations that we expect to observe in the hippocampus. One common way to represent the responses of recorded individual neurons is to plot their place fields. In Figure 4b we show the place fields (averaged over training epochs) of 36 randomly selected cells of the second layer of the network. Their fields have been measured during the simulated exploration along trajectories sampled from a distribution illustrated in Figure 4a. We observe a number of cells with localized place fields which are similar to those of classical place cells. What is most striking, however, is that the responses are highly diverse, which is typically what is observed in the hippocampus, and in particular in the dentate gyrus (8, 42).

**Fig. 4.**
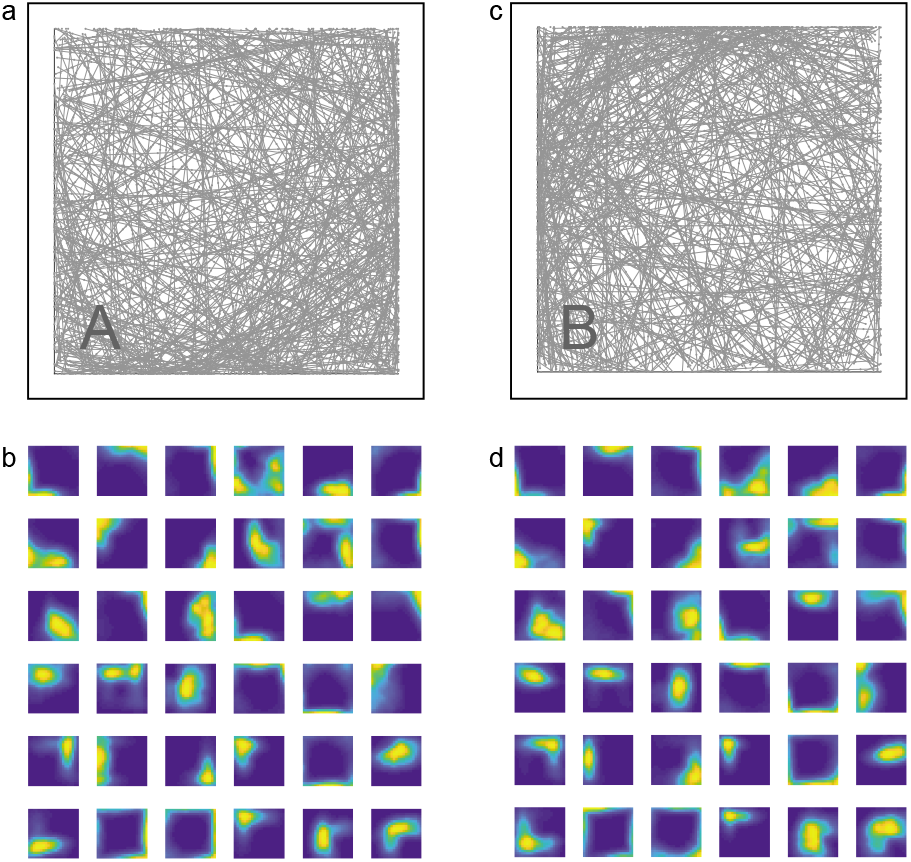
(a,b): Trajectories of a simulated animal in an open arena (exploration statistics A; top), and the spatial tuning profiles emerging from training the auto-encoder network on an artificial sensory input corresponding to these trajectories (bottom) for 36 neurons randomly selected from the second (dentate gyrus-like) layer of the model. We find a very heterogeneous set of spatial tuning profiles, some consistent with simple place cells, some exhibiting multiple place fields and some that look more like boundary cells. The statistics of this diverse set of responses appear to be consistent with calcium imaging data from the dentate (8). (c,d): Same as on the left, but for a set of trajectories with a slightly different exploration bias (exploration statistics B). Half of the trajectories on both sides have the same statistics, and are drawn from an isotropic distribution of initial positions. The other half of the trajectories are drawn from different distributions with initial positions biased towards the lower right and upper left corners, in the panels on the left and right, respectively. As a result of this, the two sets of place fields that correspond to exploration statistics A and B are slightly different.

It is interesting to consider the place fields generated by a random encoder (the connections between the first and the second layer are random). They are shown in Figure S1: the fields are substantially different and more noisy than in the case in which the weights are learned.

### The instability of place fields reflects their history dependence

Neurons with spatial tuning properties can be obtained in various ways, and while the fact that we found cells with spatial tuning in our model is reassuring, it was not unexpected given that the inputs contain information about the position of the animal. Slightly more non-trivial is the fact that the spatial tuning properties of these units are consistent with those of place cells exhibiting a small number of well-localized fields, even though the inputs consist of highly mixed representations of the spatial variables. If the inputs were provided by cells which themselves had smoothly localized spatial tuning profiles, random connectivity in combination with sparsification of the neural activity would be sufficient to achieve this (see e.g. (51)). However, for highly mixed inputs obtaining spatial tuning properties resembling those of place cells requires some learning of the weights in addition to a penalty enforcing sparse coding.

An even more interesting aspect of the responses of these cells is their dependence on recent experiences. According to our model, the neural representations in the hippocampus are continuously updated during exploration through ongoing synaptic plasticity. This means that the neural fields can be rather unstable. We illustrate this in Figure 4d where we show the place fields of the same 36 cells resulting from other epochs of the same simulation of the network during which we train on trajectories with different statistics, shown in Figure 4c. To enhance the effect of history dependence, we biased the exploration statistics in two different ways: in Figure 4a the simulated animal tended to visit the bottom right corner more often (exploration statistics A), whereas in Figure 4c the animal prefers the top left corner (exploration statistics B). Many of the neural fields are similar in the two cases, but there are clear differences reflecting the exploration bias. Due to the ongoing plasticity, these differences remain even after many training sessions with both types of exploration statistics (i.e., they don’t stem from the initial formation of the place fields during early exposures to the environment, and in fact we excluded these early sessions when computing the place field maps).

In Figure 5a we show the changes in the fields and quantify them in Figure 5b. We compute the average normalized overlap between place fields in a simulation during which the animal experiences explorations statistics A and B repeatedly in a random order (while training the auto-encoder). The average overlap between fields estimated in different sessions with the same exploration statistics is less than 0.7, and even smaller when sessions with different statistics are considered (comparing A and B sessions). This indicates that the response properties of individual neurons are continuously modified, and that they reflect the recent exploration statistics. The relatively small overlap in the case of the same exploration statistics is partly due to the stochasticity of the algorithm used to determine the synaptic weights (which includes sampling a new set of random trajectories for every session), and partly due to the dependence of the fields on the exploration statistics of the previous session (see below). Note that the overlaps of the place fields of different sessions only decay very slowly as a function of the time interval elapsed between them. This would not be the case during the initial phase of rapid learning (excluded from these plots) that occurs during the first few sessions after initializing the model with random weights (which could be thought of as a completely naive animal that has never experienced any similar environment).

**Fig. 5.**
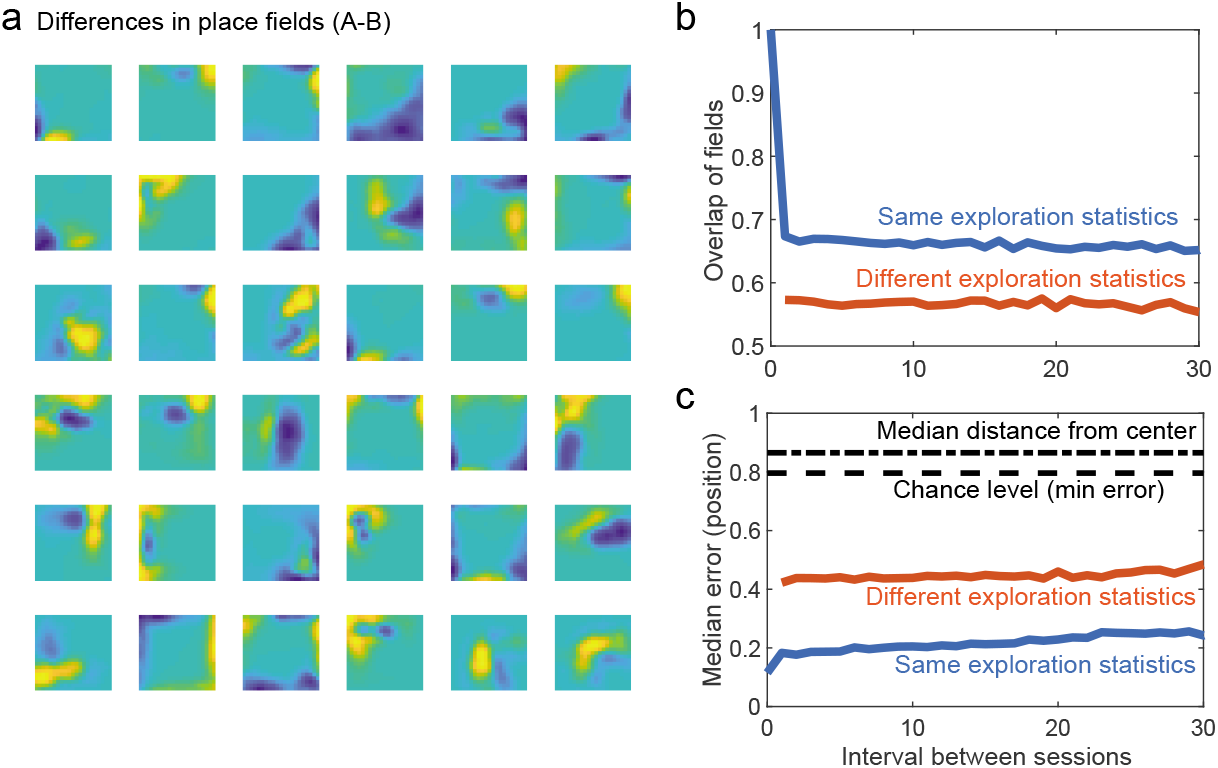
(a): Maps of differences of average place fields between A and B sessions, in a simulated experiment in which the animal experiences a random sequence of the two types of sessions with different exploration statistics (as in Fig. 4). (b): Normalized overlap between the place fields of two sessions with the same (blue) or with different (red) statistics as a function of the time interval between sessions. The overlap is larger in the former case, and stays rather high even for long time intervals between sessions, indicating relative long-term stability despite short-term fluctuations. (c): Median decoding error for position from simple regression predictors for the x and y coordinates of the animal. Position can be predicted more accurately if the decoder was trained on the same type of exploration statistics as in the session used for testing, but even for different statistics this works significantly better than chance level. The decoding error grows only slowly with the time interval between training and test sessions.

Despite the continual modifications of the fields, it is still possible to decode the position of the simulated animal, as shown in Figure 5c. We trained a linear regression decoder to predict the *x* and *y* coordinates of the animal from the second layer neural representations. The decoder is trained on the data of one session and tested on the data of a different session, as in (9). The median error is plotted as a function of the interval between these two sessions (expressed in number of sessions). The decoder is more accurate when the exploration statistics are the same, but it is still significantly better than chance (dashed lines) if they are not. Despite the instability of the fields, it is still possible to decode the position of the animal with reasonably high accuracy. This is similar to what has been observed in (9), though the statistics of the field modifications is probably different in that experiment (in which some cells respond with significant spatial tuning only in a subset of sessions; see also (52)). These differences might be due to the simplicity of our model, the fact that we are considering a 2D arena rather than a 1D track, and potentially the way the activity is recorded in the experiment. However, the model captures the basic observation that it is possible to decode position despite the relative instability of the fields.

It is worth noting that the between session variability we are quantifying here depends on a choice of parameters of the algorithm used to train the model. In our simulations, the length of an individual training session with a given exploration statistics determines the level of stability of the place fields (analogously, we can consider changing the learning rate of the animal, which plays a similar role). Furthermore, in a real experiment the within session fluctuations of the place cell responses may of course be larger than the variability due to synaptic plasticity, because of noise or additional variables encoded in the neural activity that we have not modelled here.

### History effects and the ability to decode the recent past

As already discussed, the instability of the fields is compatible with several experimental observations (see e.g. (9, 10)). However, here we propose a novel interpretation of these fluctuations that can be tested in experiments: they reflect the recent history of experiences of the animal, and therefore any bias in the exploration statistics or any other events that are represented in the input to the hippocampus should affect the neuronal responses. This means that by studying the fluctuations of the neural responses, we should be able to decode at least some information about the recent history of the animal’s sensory experiences. This is a specific prediction that can be tested if a sufficient number of neurons are simultaneously recorded for a long enough time period.

In simulations it is indeed possible to decode the recent history by reading out the fluctuations of the firing fields, but taking this approach literally one first has to construct place field maps, which requires knowledge of the position of the animal. Here we show that one can also decode some information about the previous exposure to the environment that the animal experienced directly from the neural activity patterns elicited in the current session (without requiring additional spatial information, even though much of it is of course contained in the neural representations).

We consider a random sequence of biased exploration sessions like those shown in Figure 4a,c (corresponding to a random binary string of A and B type sessions). Note that in order to ensure that discriminating A and B is sufficiently challenging, the environment, and therefore also the sensory input the animal receives given its position and head direction, is the same in A and B sessions. (If there were different landmarks in A and B sessions, that would only make the discrimination task easier). At the end of each session we estimate the place fields of the neurons in the second layer and as shown in Figures 4b,d and 5a the resulting fields depend on the exploration bias of that session. Interestingly, they also depend on the bias in the previous session: if an A session is preceded by another A, the fields (evaluated during the latter session) are different from the case in which an A session is preceded by B. We plot these differences between the fields in Figure 6a. Similarly, we show the differences in place fields (in a B session) between the case in which B is preceded by A and the case in which B is preceded by another B in Figure 6b. These differences are relatively small, but consistent enough that it is possible to train a decoder to read them out and report successfully whether the current session was preceded by A or B. In Figure 6c we show that even without first computing place fields we can train linear classifiers directly on the second layer neural representations to decode not just whether the current session, but also whether the previous session was of type A or B. While the performance of these classifiers is far from perfect when predictions are made based on a single activity vector, they can achieve very high accuracy when combining the predictions from many neural representations using a majority vote. These simple simulations illustrate one possible experiment that can be performed to test some of the central ideas of our theory.

**Fig. 6.**
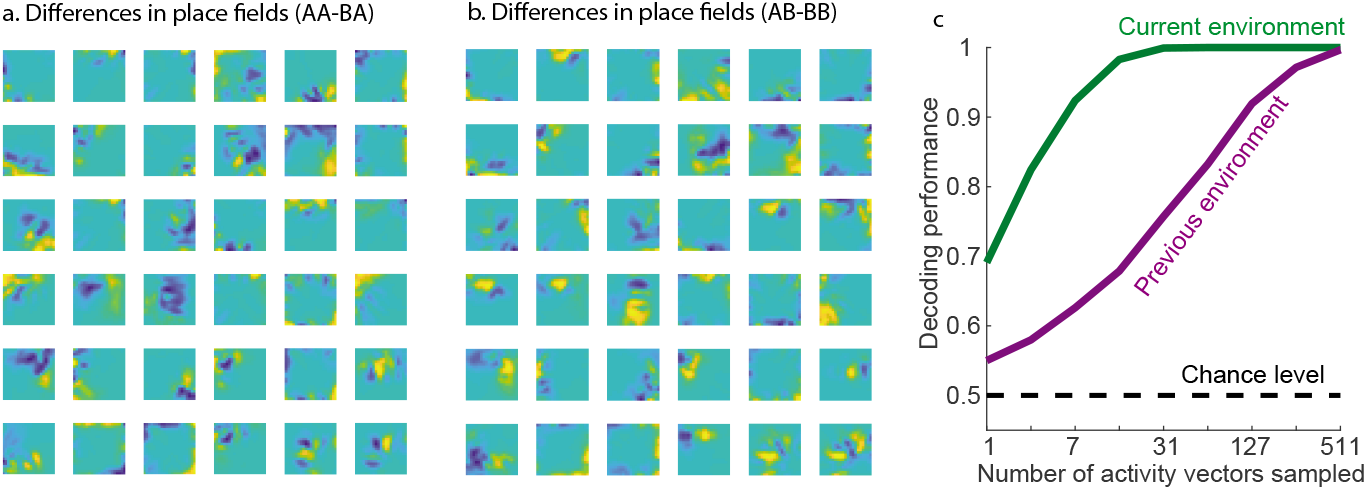
(a): Difference maps of average place fields in A sessions between the cases when the previous session was A versus B (i.e., sequences AA-BA). (b): Similar difference maps for B sessions (corresponding to sequences AB-BB). Note that these differences are more subtle than those between A and B shown in Fig. 5a. (c): To demonstrate that the fluctuations of the previous two panels are not just noise, but reliably capture history-dependent information, we show that one can decode from the neural (DG) representations of the simulated animal exploring an environment not just the statistics of the current session (i.e., A versus B; green), but also the statistics of the previous session it experienced (purple). We decode using simple maximum margin linear classifiers in combination with boosting, and report the resulting performance as a function of the number of neural representations (snapshots of the second layer activity in the current session) used for decoding. While the performance is only slightly above chance level when decoding from a single snapshot of the neural activity, a linear classifier can almost perfectly discriminate A and B sessions when combining together the predictions of the trained classifier for many such activity patterns, by taking a simple majority vote of the predicted labels. Crucially, also the decoder for the statistics of the previous session only uses activity patterns from the current session.

## Discussion

Preprocessing correlated patterns can greatly increase memory capacity. Extracting the uncorrelated components of the sensory experiences to be memorized (or more generally, the components that are truly independent from previously stored inputs) would enable the memory system to store only the information that is not already in memory. Any similarity with previous experiences can be exploited to reduce the amount of information for each input that actually needs to be stored (because it is truly novel), which decreases the amount of synaptic resources required. We proposed that the hippocampus plays an important role in this process of compressing memories, and we presented a simple neural network model that illustrates the main ideas of memory compression. This model provides a novel interpretation of the observations of numerous experiments on navigation conducted on rodents. These experiments show that the recorded neural activity in the hippocampus encodes the position of the animal, suggesting that the hippocampus plays an important role in navigation. However, there is an ongoing debate on whether the hippocampus is actually needed to navigate in familiar environments (see e.g. (53–56)). Several studies indicate that the hippocampus is important primarily in situations of navigation that require the formation of new memories (57).

Our proposal, similarly to what has been suggested by Eichenbaum (4, 57), is that the hippocampus is a general memory device used for compressing correlated inputs into efficiently storable episodic memories, and that it is only due to the nature of the navigation experiments that many investigators were led to put so much emphasis on the role of the rodent hippocampus in encoding the position of the animal. We showed that because the sensory experiences during navigation are highly correlated (for similar locations of the animal), a simple network compressing such inputs can reproduce the response properties of typical hippocampal neurons.

In our model the neural representations reflect not just current inputs, but also recent memories, since the synaptic weights are continuously updated, which would explain at least some of the observed high variability of the neuronal responses (see e.g. (10)). Moreover, our model is also in line with the elevated sensitivity of the neuronal responses to any change in the environment (e.g. the delivery of reward (12, 58), or the manipulation of landmarks (59, 60)). This phenomenon is often described as “remapping” (61) and is widely observed in experiments. In (62) the authors suggest that it might be important for memory. According to our interpretation, the hippocampus would encode any memory, not just those that are related to navigation in physical space. Hence, the presence of an item or the delivery of reward at a particular location, which often constitute salient episodes, would certainly alter the neuronal responses, as observed in experiments (see e.g. (12, 58, 60, 63)). Moreover, any structured sensory experience involving correlated inputs parameterized by some external variable would also be reflected in the neural representations in the hippocampus, as in the case of auditory stimuli studied in (23). From this point of view it is not surprising to observe hippocampal neurons develop tuning curves in the space of other external variables, such as e.g. in frequency space.

One of the testable predictions of our model is that the neuronal responses should be affected by the recent history, to the point that the fluctuations of the firing fields should contain decodable information about the recent exploration statistics of the animal. The observation of this type of history effects would be only the first step to corroborate the validity of the model. Indeed, the observation of history effects does not guarantee that the hippocampus is compressing sensory inputs. To confirm that the role of the hippocampus in memory compression is as suggested by the model we would need additional experiments in which the statistics of the sensory experiences are structured, and for example organized as in the ultrametric case that we discussed at the beginning of the article. Comparing the neural representations before and after learning should reveal whether the changes are compatible with a compression process (e.g. by looking at the geometry of the representations, as in Figure 2 and its dependence on the compressibility of the environment). Moreover, it is important to stress that history effects are not a unique feature of our model. During learning, it is likely that any model would exhibit history effects. However, if the hippocampus was designed to encode only the position of the animal as accurately as possible, it would appear unlikely that in a stationary environment the representations would keep changing substantially once the environment is sufficiently familiar and the position of the animal can be decoded from the neural activity. Our model predicts that even in situations in which position is strongly encoded, we should observe continual modifications that reflect the recent history of sensory experiences.

There are other recent models that focus on the navigational role of brain areas like entorhinal cortex, and nevertheless predict that spatial representations should be continuously updated. In particular, in (64, 65) the authors assume the responses of neurons (grid cells) in entorhinal cortex are modified whenever a landmark is encountered to correct for the errors that accumulate in an intrinsically noisy process of path integration, which introduces history (path) dependence of their firing fields. Some of these predictions have already been successfully tested in experiments (64, 65). In the future it will be interesting to study whether our model of the hippocampus is compatible with these observations in entorhinal cortex. It is worth noting that the models proposed in (64, 65) are constructed to explicitly solve navigation problems, whereas our main hypothesis is that the hippocampus cares primarily about efficient memory storage, and place cells, which could be used to navigate, emerge naturally as a consequence of a memory compression process. Our model can be applied to any task that requires memory, whether it involves navigation or not. The models of (64, 65) can probably be generalized to other tasks, but they might still require a structure in the input space that is similar to the one studied for real space navigation.

There is also another category of recent theoretical work that emphasizes the general role of the hippocampus in learning and memory (see e.g. (66–73)) at the expense of the specific role it plays in navigation. These works mostly focus on the encoding of temporal sequences with one notable exception (73) and some assume that the hippocampus tries to store only the information that is relevant for predicting the next state of the environment, while others postulate that the goal of the hippocampus is to represent a whole probability distribution of future locations (conditioned on the current position of the animal). These predictions are then used to drive reinforcement learning. In our case, we considered for simplicity only the compression of the instantaneous sensory experiences. We focused on the lower level question of how spatially modulated cells may arise mechanistically, and which of their features may be explained without postulating any higher cognitive goals other than simply remembering the past. The model can easily be extended to deal with simple temporal correlations, for example by replacing the input layer with a recurrent network. Even in the case of random connectivity, the network activity would then contain information about the temporal sequence of recent sensory experiences (74). For longer timescales and more complex temporal correlations a different mechanism would be required (see e.g. (75, 76)), which might involve also synaptic plasticity on multiple timescales (17, 77). In all these cases the instantaneous input of the autoencoder contains information about a recent temporal sequence. The downstream autoencoder can then be modeled exactly in the same way as we modeled it here. The development of new models of the input that can deal with temporal correlations will be one of the priorities of our future research.

Finally, in our work we focused on the role of the hippocampus in compressing a stream of correlated sensory experiences: the correlations that we considered are only those present in the input patterns, which we assumed represent the sensory experiences of the animal. However, it will be important in the future to consider also the correlations between the sensory input and long-term memory, which probably resides in the cortex (see e.g. (78, 79)). Hippocampus is anatomically part of a loop that involves the cortex, and this loop is important for at least two reasons: the first one is that the hippocampus can contribute greatly to organizing long-term memories stored in the cortex, and it might be important for the process of abstraction that underlies the creation of schema. The second one is that the hippocampus should take into account also the similarities between the current episode and all the memories already stored in the cortex. Even more importantly, the hippocampus should be able to use the abstract information that might be stored in the long-term memory. A recent model (72) addressed these two issues by introducing a mechanism that maps specific episodes to low-dimensional structures that are encoded in the cortex. The model also assumes that the hippocampus plays an important role in the formation of these cortical representations. Moreover, the authors show that such a model leads to the formation of place cells in the hippocampus and grid cells in entorhinal cortex. Our model focused more the role of the hippocampus in episodic memory and it considers for simplicity only spatial correlations between sensory experiences. In (72) instead the authors assume that the instantaneous sensory experiences are completely random and uncorrelated and the only correlations that are considered are temporal. Incorporating into our model a loop with the cortex in a way that is similar to what has been proposed in (72) would probably lead to even more efficient ways of storing episodic memories and will be considered in future studies.

## Materials and Methods

### Memorizing ultrametric patterns

The patterns to be memorized are organized as in an ultrametric tree (see e.g. (25–28)). We can consider the case of *p* ancestors, each with *k* descendants. The ancestors are random dense binary patterns, and the descendants are obtained from the ancestors by resampling (i.e., randomly picking a new binary value for) the activity of each neuron with probability 1 – *γ*, such that their normalized overlap with their ancestor is equal to *γ*. For *γ* = 1 the descendants are identical to the ancestors, whereas for *γ* = 0 they are completely independent.

To store the memories given by all the descendant patterns we can use two networks of *N* neurons each, one for storing the ancestors, and the other to store the differences between ancestors and descendants. The first network has connectivity *c*_1_ (where *c*_1_ is the number of incoming connections per neuron) and the second one has connectivity *c*_2_.

To store *p* ancestors in the first network one must have

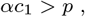

where *α* is the coefficient of the memory capacity (we are assuming that the number of patterns that can be stored is proportional to the connectivity per neuron).

In the second network the coding level *f* is proportional to 1 – *γ* (for *γ* close to one) and the constraint on *c*_2_ will approximately take the form

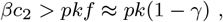

with *β* being another capacity coefficient of order one, which does not depend strongly on the parameters *N*, *f* or *γ*.

The minimum number of synapses (in both networks combined) that are needed is therefore

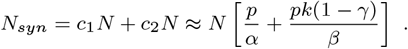

If we denote by *P* = *pk* the total number of patterns, we find

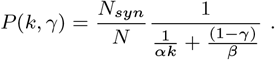

One case in which the patterns are all uncorrelated is when *k* = 1 and *γ* = 0. We define this case as *P*_uncorr_ = *P*(1,0). In Fig. 1b we plot *P*(*k, γ*)/*P*_uncorr_ as a function of *k* and *γ*, which gives the relative advantage of using the compression strategy when the patterns are correlated, as opposed to storing incompressible patterns. We used *α* = 0.1 and *β* = 0.025.

### Simulations of the compression of ultrametric patterns

The simulated network is illustrated in Figure 2a. The ultrametric patterns described in the previous section are imposed on the input layer (EC in the figure). To generate the compressed representations of the second layer (DG) we trained the auto-encoder of Figure 1a using the standard matlab algorithm `trainAutoencoder`. We used the default activation function which is a logistic sigmoid. No noise was added to the ultrametric patterns. As the number of neurons in the second layer is larger than in the input layer (*N* = 300 for the input layer, *N* = 600 for the intermediate layer), we compressed the neural representations by using the sparseness regularization option. In particular, we chose the following parameters: `‘SparsityProportion’ = fsparse, ‘SparsityRegularization’ = 5, ‘L2WeightRegularization’` = 0.00001, where fsparse was the desired level of sparseness. The maximum number of training epochs was 600. Once we obtained the sparse hidden layer activity patterns for all ultrametric patterns we binarized the representations by setting the neuronal activity to 0 or 1 depending on whether it was below or above the threshold Θ = 0.5. The results do not depend much on the choice of the threshold because the distribution of the activities in the second layer was usually markedly bimodal. The coding level (sparseness) of the binarized representations was close to the one specified by the parameter `fsparse`.

The geometry of the auto-encoder representations (in particular their correlation structure) was compared in Figure 2b,c to the geometry of random representations, which were obtained using random projections of the same ultrametric input patterns (see e.g. (39–41)). These representations were constructed by choosing the weights of the connections between the input and the second layer randomly. The weights were independent Gaussian variables with zero mean and unit variance. The results do not depend much on the specific choice of the distribution of the weights. We then determined the activity of the neurons in the intermediate layer by thresholding the total synaptic current. The threshold was chosen to reproduce the same sparseness of the binarized second layer representations as for the trained auto-encoder.

In Figures 2b,c we computed the overlaps between the neural representations for the input, for the second layer of the autoencoder and for the second layer of the random projection network. Different points correspond to different branching ratios (*k* = 10, 20, 30, 60, 75, 100, 150 and 300). The total number of ultrametric patterns was kept constant (*P* = 300) by decreasing the number of ancestors when the branching ratio increased. In these figures the sparseness level was `fsparse` =0.1 and *γ* = 0.6.

In Figures 2d,e we estimated the memory capacity. We progressively increased the number of memories by changing the number of ancestors. The branching ratio was fixed (*k* = 2 for one curve in panel d, and *k* = 20 for all the other curves). After training the auto-encoder, we used the binarized representations to reconstruct the input. To obtain the weights between the second layer and the layer containing the reconstructed representations we used the matlab algorithm `fitclinear`. We then tested this memory system by presenting to the input memory cues which were noisy versions of the stored memories (the average overlap between the cues and the original memories was 0.8). A memory was considered retrievable if the reconstructed memory had an overlap of at least 0.9 with the stored memory. In Figure 2d,e we plotted the number of memories successfully retrieved under these conditions.

We adopted a similar procedure to estimate the number of retrieved memories also for the network with random projections and for a direct readout from the input. In the latter case we reconstructed the activity of each input neuron from the other neurons (i.e., the reconstructed neuron did not have access to the corresponding input neuron).

The memory capacity depends on the compressibility of the neural representations (branching ratio *k* and similarity between descendants and ancestors *γ*) and on the coding level *f*. *f* was chosen to be optimal among *f* = 0.05, 0.075,0.1, 0.125,0.15. The other parameters were: *γ* = 0.6, 0.2, input layer size: *N* = 300; second layer size: *N* = 600.

### Simulations of input compression in navigational experiments

The sensory input of the simulated animal can be modeled in various ways, and we have experimented with different forms of e.g. simulated visual inputs. Here we describe the simplest version used to generate the figures above, which involves specifying three variables for every straight line excursion of the animal through the environment. The first variable is the starting position along the wall as given by an angle *ϕ* between a fixed direction (say East) and the vector pointing towards the starting position (at the wall) from the center of the arena. The second one is the direction of motion, expressed as an angle *θ* measured with respect to (the vector orthogonal to) the starting wall, and the third is the distance *d* travelled since the last wall contact. Note that because we focus on straight line trajectories, these three measurements uniquely determine the animal’s position in the environment, though each location corresponds to many different triplets of values for these three variables. In order to put these inputs on an equal footing, we take the sines and cosines of these variables (after converting the distance *d* into an angle by multiplying by 2*π* and dividing by its maximum value, which is 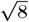 in a square arena with x and y coordinates valued in [−1, 1]). Then we pass these variables through a random projection *W_inp_* and an element-wise step function *H* (which expands the dimensionality by mixing the trigonometric spatial variables non-linearly) to obtain binary representations for the input to our network:

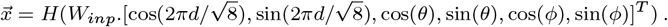

Note that the random projection *W_inp_* used to compute these input patterns 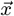 is different from the random projection between the input and second layer we use in the control model to compare to the model with learned feed-forward weights.

These input representations 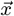 are then sampled along a set of trajectories (500 per session) constructed as follows. First we sample an initial position in the form of an angle *ϕ* at the center of the arena corresponding to the vector pointing towards the starting point along the wall. For half the trajectories this angle is sampled uniformly from the interval [0, 2*π*], while for the other half *ϕ* is given by a sample from the uniform distribution on [0, *π*] plus a bias term *bπ*/2 + 3*π*/4. Here *b* is a binary variable that equals 1 in A sessions and −1 in B sessions. A heading direction *θ* is then sampled uniformly in [−*π*/2, *π*/2], with *θ* = 0 corresponding to motion orthogonal to the starting wall, and finally the animal moves from its initial position in this direction. The distance *d* from the starting point increases along the excursion into the environment, and we record an input vector 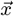 for each distance increment of 0.1 until the animal encounters another wall, at which point the trajectory is complete and we repeat the procedure for the next one.

The set of correlated input patterns 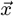 constructed in this manner for one session in the environment, which we will use as our putative EC representations, is analogous to the set of ultrametric memory patterns discussed above. Again, we will train a neural network model (in particular the synaptic weights between EC and DG of the animal) to compress them in order to store these sensory inputs efficiently. After each session we use the input patterns recorded while the animal was exploring the environment to train an auto-encoder with a sigmoidal non-linearity in the hidden layer (as well as in the readout layer, with an architecture as shown in Fig. 2a) to reconstruct these patterns despite a sparsity penalty enforcing a bottleneck in the DG layer. The size of the network is the same as above, and the parameters used for training are `‘MaxEpochs’ = 10, ‘SparsityProportion’ = 0.03, ‘SparsityRegularization’ = 1 and ‘L2WeightRegularization’ = 0`. Note that for simplicity the training occurs offline (i.e., in batch mode, after the session) using back-propagation as implemented in the matlab routine `trainAutoencoder`. Thus the ordering of the input patterns and the fact that they were recorded in a particular temporal sequence along the sampled trajectories play no role (i.e., we would expect the same results if we randomly permuted the order of the patterns within the set of inputs). We repeat this training procedure for 60 sessions with independently sampled sets of trajectories (and therefore input patterns), choosing a random exploration bias *b* for each session.

To assess the spatial tuning properties of the resulting DG representations, we construct maps (of 21 by 21 bins) of the cumulative activity of individual hidden-layer units, and normalize them by the total occupancy of each spatial bin. Before plotting the resulting maps as in Fig. 4, we smooth them by convolving with a Gaussian filter of width one bin. The place field maps shown in this and subsequent figures are averages over all the sessions of a given type, but excluding the transient learning period that occurs during the first 20 sessions after initializing the network with random weights.

In order to study the memory capacity of the network, we remove the (non-biological) third layer of the auto-encoder, and feed the second-layer representations obtained from the trained auto-encoder into a CA3-like auto-associative memory module with recurrent connections. The architecture of the network model is as shown in Fig. 3b. We assume for simplicity one-to-one (identity) connections between DG and CA3, but again binarize the DG representations with a threshold of Θ = 0.5. The resulting coding level of these binary representations was significantly smaller than the target value 0.03 during the initial learning, but only slightly smaller than the target value in the sessions after this transient period. We checked that in these sessions the input patterns can be accurately reconstructed both from the continuous-valued hidden layer representations of the trained auto-encoder and from their binarized versions (with only a modest reduction in accuracy).

The auto-associative network stores the binary representations obtained from the trained auto-encoder by one-shot learning using a normalized covariance rule to update the recurrent connections. More precisely, in order to avoid overloading the memory network (which only has *N* = 600 neurons) we distribute the hidden layer representations corresponding to one session across four different copies of this auto-associative network (storing each activity vector in only one of them), and report the average memory capacity of these networks in Fig. 3c. In order to assess this memory capacity, we initialize the neural activity with noisy versions of the stored activity patterns, and then iteratively (and simultaneously) update the states of the binary neurons by thresholding their input currents (sums of mean-subtracted presynaptic activities times weights). The (approximately) optimal thresholds are determined numerically by a simple line search maximizing the number of correctly retrieved patterns. We count a memory as successfully recalled if the binary network states obtained after 10 iterations of the network retrieval dynamics has a normalized overlap larger than 0.8 with the corresponding (compressed) binary representation that was originally stored. The noisy cues used as initial states for retrieval are obtained by randomly flipping bits of the stored patterns until their normalized overlap is lower than 0.7. This ensures that the network has to reconstruct information not present in the cue for a memory to be considered correctly recalled. We again compare this memory performance to that of a network of the same architecture and coding levels, but random (instead of learned) synaptic weights between the input and second layers.

Decoding the statistics of the preceding session (specifically, the binary variable *b* that determines the exploration bias in our case) from the compressed activity patterns recorded during the current session can be done in variety of ways. One of the simplest approaches is to train a linear classifier (again we use the support vector machine algorithm `fitclinear`) to discriminate activity patterns sampled at different times from sessions preceded by an A session from those preceded by a B session, and evaluating the test performance in a cross-validated fashion on held out patterns. This leads to an average performance (the classification accuracy is the average over all held out patterns) that is already above chance level, as shown in the left-most data point in Figure 6c (purple). This is the average performance one would get by reading out a randomly chosen pattern of activity at a particular time. We can further enhance the performance by combining together the predictions of the classifier for multiple test activity patterns that correspond to representations at different times. To do so, we construct two equally sized test sets by sampling a given number of held out patterns from sessions that were preceded by an A versus a B session. Then we assess the ability of the decoder to discriminate between these two test sets by simply counting the number of correct predictions on each of these test trials, and calling the overall classification correct if it is larger than the number of wrong predictions on the test trials. For comparison, we also show the cross-validated classification performance for the exploration bias *b* of the current session (green). The fact that this can also be decoded is less surprising, since the current exploration bias clearly changes the statistics of the activity patterns in the present session even in the absence of learning.

## ACKNOWLEDGMENTS

We thank A. Losonczy and J. Priestley for many useful discussions. We are grateful to D. Aronov, R. Gulli, A. Losonczy, E. Mackevicius and J. Priestley for comments on the article. This work was supported by DARPA L2M, NSF’s NeuroNex program award DBI-1707398, the Gatsby Charitable Foundation, the Simons Foundation, and the Swartz Foundation.

---

**Fig. S1.**
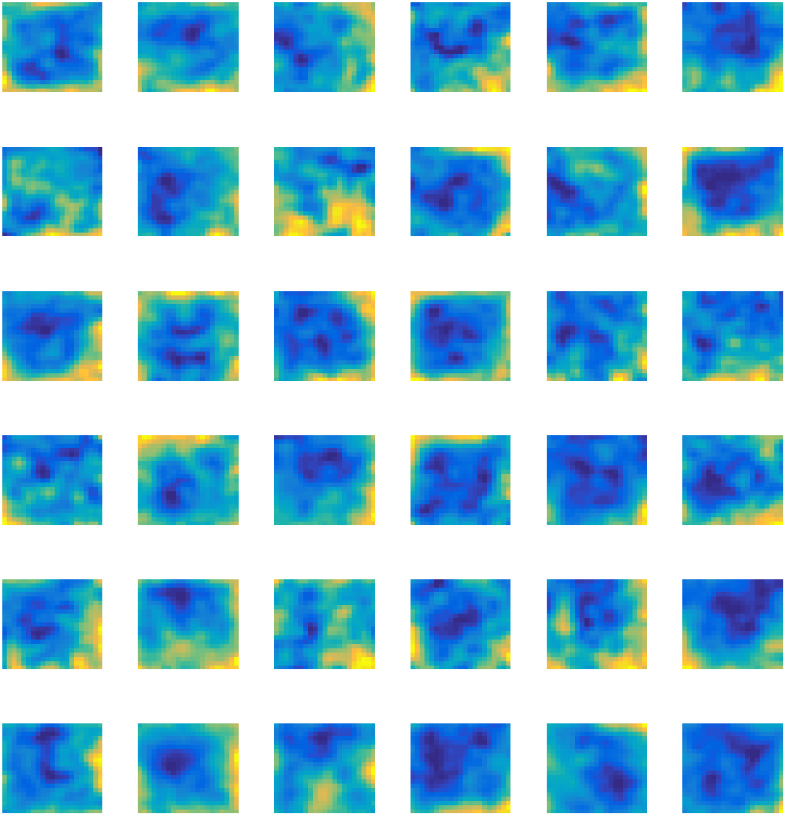
Spatial activity maps of the second layer neural activities for a network of the same architecture as in Fig. 4b and d, but with random (instead of learned) synaptic connections between the first and second layers. While there is still some residual spatial selectivity (by chance), the activity maps are not as coherent and much less reminiscent of experimentally observed place fields.

